# The APOL1 variant p.N264K blocks ion flow by occluding a pore at the cell surface

**DOI:** 10.1101/2025.11.05.686714

**Authors:** Verena Hoeffken, Lara Console, Niklas Nelde, Hermann Pavenstädt, Cesare Indiveri, Thomas Weide

**Author notes:** These authors contributed equally (Verena Hoeffken, Lara Console, Cesare Indiveri, Thomas Weide).

## Abstract

The APOL1 gene variants G1 and G2 are associated with an increased risk of APOL1-mediated kidney disease. A recently identified variant, p.N264K (M1), mitigates this risk of renal damage by abolishing APOL1-G2’s associated cytotoxicity. However, the molecular and structural basis of this protective effect remains incompletely understood.

In this study, we first show that both the cytotoxic G2 and the non-toxic M1-G2 exhibit similar intracellular localization, surface expression, and turnover kinetics. Moreover, N-glycosylation assays indicated no differences in topology, and 3D models demonstrated that both cytotoxic G2 and non-toxic APOL1 M1-G2 span the membrane four times, forming a potential ion channel.

Interestingly, molecular dynamics analyses further revealed that in M1-G2, the lysine at position 264 occludes this channel, thereby preventing ion pore activity of APOL1. These findings provide, for the first time, a mechanistic explanation for the non-toxic behavior of the APOL1 M1-G2 variant. Additional 3D analyses suggest that the C-terminal region may contribute to APOL1 multimerization, potentially influencing ion flux and cytotoxicity.

## INTRODUCTION

Humans are protected against protozoal infections by African trypanosomes, because they are inactivated by the human serum protein Apolipoprotein L1 (APOL1), except for the subspecies *Trypanosoma brucei gambiense* and *rhodesiense*, which cause human *African* sleeping sickness (*Human African trypanosomiasis, HAT*) (Vanhamme *et al*, 2003; Perez-Morga *et al*, 2005). Co-evolution between host and parasite resulted in two APOL1 variants able to restore protection against blood parasites. These variants, termed G1 and G2, are characterized by two amino acid (aa) substitutions (S342G and I384M; G1) or two deletions (del:N388Y389, G2) in the C-terminus of APOL1 (Pays *et al*, 2006, Pays *et al*, 2023). The presence of two of these alleles (G1/G1, G2/G2, G1/G2) is associated with an increased risk for a wide spectrum of kidney diseases summarized as APOL1-mediated kidney diseases (AMKD)(Tzur *et al*, 2010; Genovese *et al*, 2010; Friedman & Pollak, 2021; Daneshpajouhnejad *et al*, 2022).

In 2013, the characterization of an atypical *T. b. gambiense* infection demonstrated the existence of a further APOL1 variant, p.N264K (hereafter M1)(Gupta *et al*, 2023). In homozygous carriers, this substitution may reduce trypanolytic activity, potentially enabling trypanosomes to evade APOL1-mediated protection (Cuypers *et al*, 2016). Strikingly, recent clinical data showed that this variant is not associated with AMKD, even when combined with two APOL1 G2 alleles (Gupta *et al*, 2023).

The replacement of asparagine by lysin lies in the membrane-addressing domain (MAD), adjacent to the pore-forming domain (PFD) of APOL1. Both domains are thought to play a critical role in APOL1’s binding to membranes (Höffken *et al*, 2025b). Moreover, M1 localizes next to a putative cholesterol-binding site and is also close to a region that has undergone positive selection during evolution (Müller *et al*, 2021; Lecordier *et al*, 2023). Unlike APOL1-G2 alone, the combination of APOL1-G2 with M1 possessed no ion flux, which supports the role of APOL1 as an ion pore (Olabisi *et al*, 2016; Datta *et al*, 2024; Bruno *et al*, 2017; Giovinazzo *et al*, 2020; Schaub *et al*, 2020), and indicates that this position may represent a key site for APOL1-asscociated cytotoxicity (Hung *et al*, 2023). However, it remains unclear *why* this specific aa substitution interferes with ion transport.

In this study, we investigated this aspect in more detail by directly comparing cytotoxic APOL1-G2 with the non-toxic APOL1 M1-G2 variant. Using a combination of cell-biological and *in silico* approaches, we assessed total and surface expression, protein turnover, and membrane topology of both APOL1 variants. In addition, *AlphaFold3*-based analyses were performed to compare their 3D structures and potential dimerization or oligomerization properties.

## RESULTS

The APOL1 M1 variant occurs either with wildtype (G0) or with the G2, but never with the G1 variant (Hung *et al*, 2023; Simeone *et al*, 2025). To compare the cytotoxic G2 with the non-toxic M1-G2 variant, we generated stable HEK293T cell lines enabling Doxycycline (Dox)-dependent expression of untagged or C-terminally GFP-tagged APOL1-G2 and M1-G2 (Fig. 1A) (Müller *et al*, 2021). Western blot analyses demonstrate a clear Dox-dependent expression of both variants. Next, we used viability assays of cells expressing G2 or the M1-G2 (Fig. 1B) and observed - as expected - that the M1 variant abolishes the cytotoxic effects of APOL1-G2 (Fig. 1C). APOL1 is thought to adopt multiple conformations, including one with four transmembrane (TM) regions (Fig. 1A,D) (Gupta *et al*, 2020). Since the M1 substitution lies in the putative 3^rd^ TM of APOL1, it could alter membrane topology, reducing APOL1 TMs from four to three resulting in a relocation of the C-terminus from the *endoplasmic reticulum* (ER) lumen to the cytoplasm. To test this, we added N-glycosylation tags to APOL1-G2 and M1-G2 and analyzed glycosylation, which occurs only if the C-terminus is in the ER lumen(Müller *et al*, 2021). Using PNGase F assays as described before (Müller *et al*, 2021), our data show no evidence of altered topology, indicating that APOL1-G2 and M1-G2 share a similar membrane orientation (Fig. 1E).

**Figure 1:**
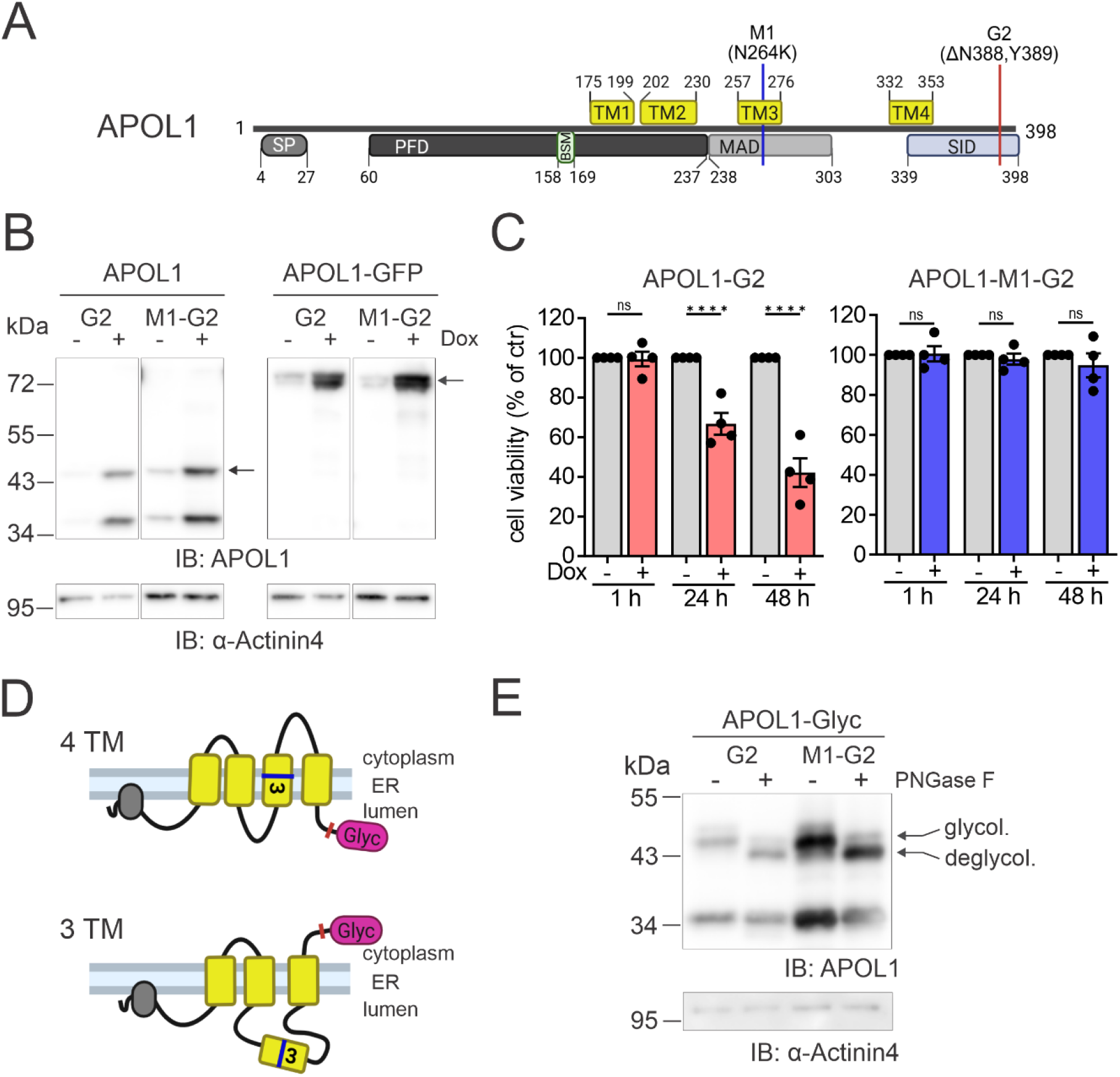
Inducible expression of the M1 variant of APOL1-G2 showed no cytotoxicity while similar orientation within the ER membrane. **(A)** Scheme of the APOL1 sequence with four transmembrane domains (TM1-4, yellow) and structural features signal peptide (SP), pore forming domain (PFD), membrane addressing domain (MAD) and SRA-interacting domain (SID) as well as the BH3 domain-like sequence motif (BSM). Variant positions M1 (N264K) and G2 (del:N388Y389) are indicated. **(B)** Western blot (WB) analysis of stable HEK293T cells verified expression of APOL1-G2 with or without the M1 variant (left) or with an additional C-terminal GFP tag (right) after induction by Doxycycline (+Dox, 125 ng/ml) for 24 h. **(C)** Cell viability assay of APOL1-G2 (left) and APOL1-M1-G2 (right) showed cytotoxicity after Dox induction over time for G2 but not for M1-G2 when compared to uninduced conditions. N = 3, n = 6; ^****^ = p <0.0001. Values shown as means with SEM. **(D)** Schematic model of possible topological orientations of APOL1-M1-G2 with four transmembrane regions (4TM) or three (3TM) inserted in the membrane. A glycosylation tag (Glyc) was added at the C-terminus to investigate APOL1’s topology. **(E)** WB analysis of APOL1-G2 and APOL1-M1-G2 proteins with or without PNGase F treatment removing glycosylation. Both variants showed a clear shift downwards after digestion indicating an identical topology with C-term facing the ER lumen.

We next examined whether the distinct properties of APOL1-G2 and M1-G2 result from differences in intracellular localization, as particularly reduced plasma membrane (PM) levels of M1-G2 could explain the decreased ion transport. IF analyses of stable cell lines expressing GFP-tagged APOL1-G2 and M1-G2 showed that both variants localize mainly to intracellular, likely ER-associated pools, with significant signals also detected at the PM (Fig. 2A, Fig. EV1). FC with the same cell lines confirmed PM-localized APOL1 pools of comparable levels as shown by transient expression experiments (Fig. 2B, Fig. EV2). Since differences in APOL1 ion activity at the PM may result from faster degradation (Höffken *et al*, 2025a) of M1-G2, we compared the turnover of both variants and found, however, that both variants exhibit similar degradation kinetics: ER-associated fractions are rapidly degraded, whereas PM-associated pools remain relatively stable (Fig. 2C, Fig. EV1). Thus, together, our data reveals no major differences between G2 and M1-G2 with respect to membrane topology, intracellular distribution, or turnover dynamics, indicating that the functional divergence between the toxic G2 and the non-toxic M1-G2 variants primarily arises from intrinsic structural differences enabling their ion channel activity.

**Figure 2:**
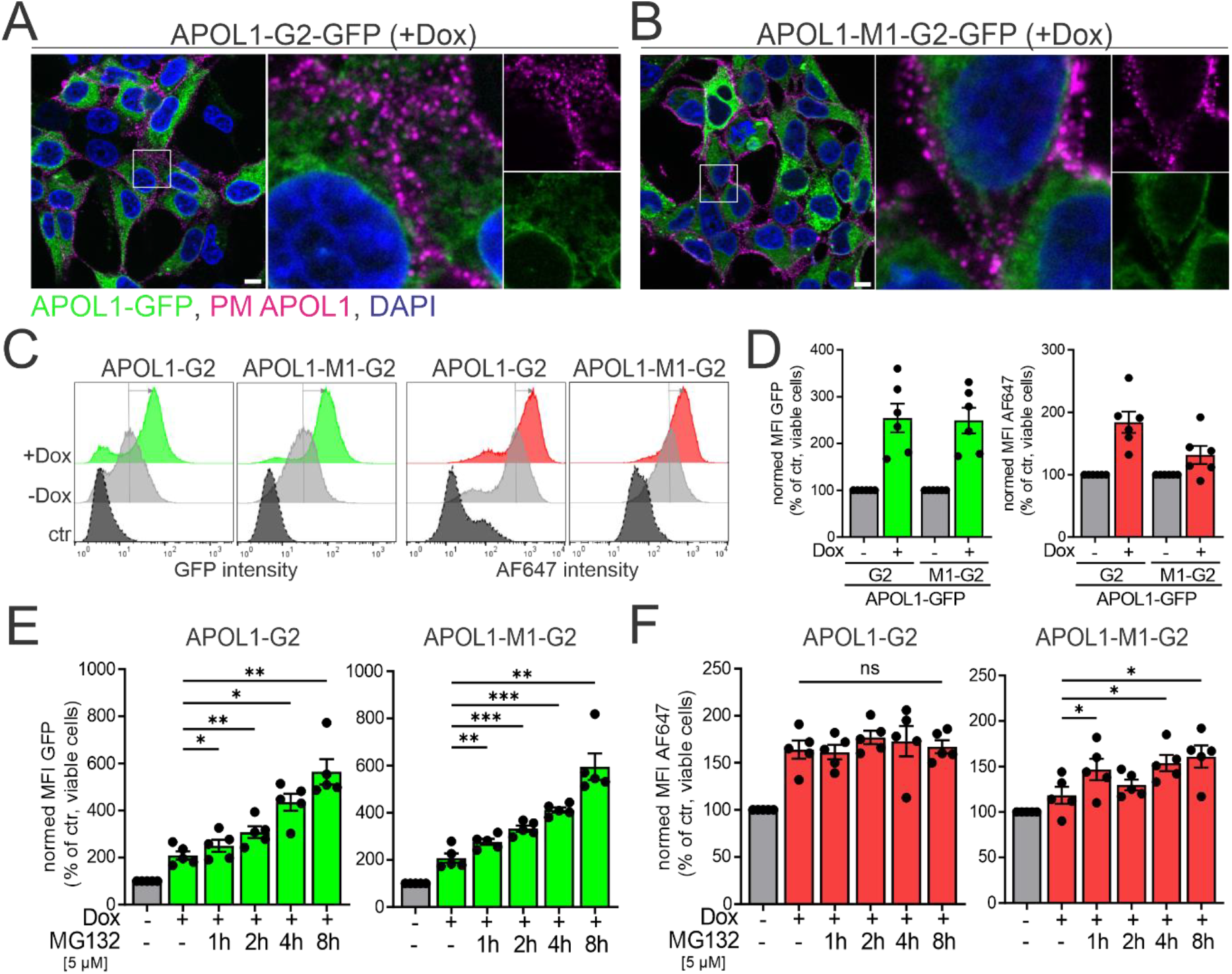
Surface-localized APOL1 pools were detected in both the M1 variant and APOL1-G2 indicating efficient transport to the PM. **(A, B)** Immunofluorescence (IF) staining of unpermeabilized HEK293T cells expressing APOL1-G2-GFP or with additional M1 variant after 24 h of Dox-induction (125 ng/ml). While GFP signal displays total APOL1 protein within the cells, specific antibody staining against APOL1 (magenta) highlights punctuated surface-localized APOL1 proteins in both cell lines. Nuclei were counterstained with DAPI. Scalebars = 10 µm. **(C)** Histogram of flow cytometric (FC) analysis of APOL1-G2-GFP and APOL1-M1-G2-GFP verified an increase in GFP signal as well as APOL1 surface signal (AF647) after 24 h of induction (+Dox, 125 ng/ml) compared to uninduced cells (-Dox) and ctr. **(D)** Normed mean fluorescence intensities (MFI) of GFP (left) and AF647 intensities (right) of FC analysis showed positive expression of APOL1-GFP after induction as well as presence of surface-localized pools for both the G2 and the M1-G2 variants. (N = 6). **(E, F)** Normed MFIs of GFP intensities representing total APOL1 pools (E) and AF647 intensities representing surface pools (F) after induction and additional inhibition of proteasomal degradation by MG132. While GFP signals increased rapidly over time for both cell lines after degradation inhibition, AF647 signal remained stable for G2 or only slightly increased for M1-G2. N = 5, ^******^ = p<0.01/0.001/0.0001. Values shown as means with SEM.

We used *AlphaFold3* and *RosettaFold* to investigate APOL1’s 3D structure (aa residues 171-360) in more detail. Both predicted a 3D structure with 4TMs (Fig. 3, Fig. EV2). Differences between the *in silico* tools likely arise from *RosettaFold*’s inability to model fatty acids that mimic the membrane’s hydrophobic environment, what *AlphaFold3* can. Therefore, the *AlphaFold3* structure was selected for molecular dynamics (MD) simulations to assess stability in a membrane-like environment, including the previously identified transmembrane region. Confidence scores from the server, such as pLDDT (>90% in several regions) and pTM (0.54), indicate a reliable model (Fig. 3A).

**Figure 3:**
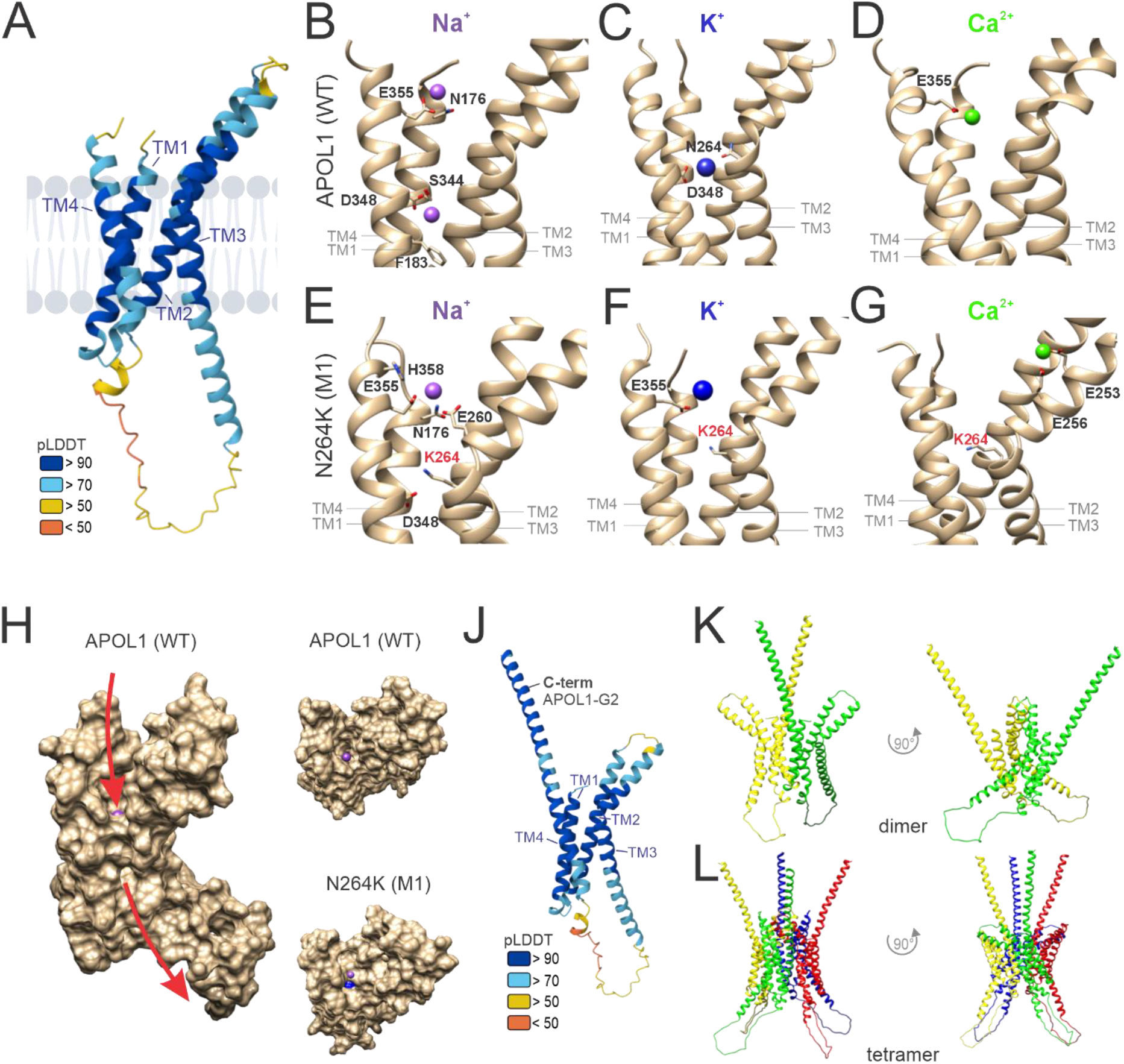
AlphaFold3 modeling of APOL1 confirmed four transmembrane regions and indicates pore blocking by M1. **(A)** The *in silico* prediction of APOL1 structure (aa 171-360) using AlphaFold3 confirmed four TM regions (TM1-4), shown with a ribbon diagram colored according to the pLDDT accuracy score. (**B - D**) Enlarged view of APOL1 WT showing residues involved in the interaction with a sodium ion (Na+, purple; B), a potassium ion (K+, blue; C) or a calcium ion (Ca2+ green; D). (**E - G**) Enlarged view of the APOL1 M1 (N264K) variant showing the central position of the K264 as well as residues involved in the interaction with a Na+ (E), K+ (blue, F) or a Ca2+ ion (green, G). (**H**) Solid surface visualization shows possible ion flux through the APOL1 WT pore, while the central lysine of the M1 variant (blue) blocks the ion path. (J) Predicted structure of TM segments with C-terminus of APOL1-G2 (aa 171-end) shown as a ribbon diagram colored according to the pLDDT accuracy score. (**K, L**) Two possible dimer and tetramer assemblies of the APOL1-G2 variant predicted by ClusPro.

During MD, Na^+^ ions frequently bound two regions within a pore-like stretch lined by four α-helices. The residues with the most contacts were D348, in the pore center, and E355 at the pore surface (Fig. 3B). Both have been previously shown to be critical for pore gating and cation selectivity (Schaub *et al*, 2020, Schaub *et al*, 2021). K^+^ ions also interacted with D348 (Fig. 3C), whereas Ca^2+^ ions bound mainly to E355 (Fig. 3D), suggesting a specificity of APOL1 for monovalent ions. MD simulations of the M1 variant revealed that Na^+^ and K^+^ interacted only with E355, as the K264 side chain occludes the pore and prevents binding to D348 (Fig. 3E, F). Moreover, the interaction with Ca^2+^ was even weaker (Fig. 3G). This behavior supports M1’s protective effect against APOL1-G2 mediated ion efflux (Fig. 3H). Including the C-terminal 38 aa in APOL1 led to low-confidence predictions from both *AlphaFold* and *RosettaFold*. In contrast, the same C-terminus with the G2 variant deletion restored *AlphaFold* confidence above the threshold, resulting in a rod-like extension following the 4^th^ transmembrane helix (Fig. 3J). Finally, we analyzed potential multimerization of APOL1-G2 using *ClusPro*. The results suggest that APOL1-G2 (as well as M1-G2) may form dimeric or tetrameric assemblies at the cell surface (Fig. 3K, L).

## DISCUSSION

Our analyses show that the toxic APOL1-G2 and the non-toxic M1-G2 variants are largely similar in terms of intracellular distribution, membrane topology, and turnover. 3D structural analyses further showed that both variants can span membranes four times, as proposed before (Müller *et al*, 2021; Schaub *et al*, 2020, Schaub *et al*, 2021; Giovinazzo *et al*, 2020; Gupta *et al*, 2024) supporting the potential formation of an APOL1 ion pore at the cell surface. However, interestingly, the 3D models also suggest that the lysine substitution in the 3^rd^ transmembrane domain (K264) blocks potential pore activity, thereby preventing the passage of monovalent cations. These findings provide a coherent mechanistic explanation for how the APOL1-M1 variant mitigates the cytotoxic effects of APOL1-G2.

Remarkably, the renal risk variants (RRVs) G1 and G2 are not located within the pore-forming α-helices. This raises the question of *how* the APOL1 C-terminus enables ion conductance in the case of APOL1 RRVs, and conversely, *how* it prevents it in the APOL1 wildtype proteins. Our results also show that APOL1-G2 can form oligomers, likely involving the “rod-like” C-terminus. Our results are consistent with previous reports showing that the residues D348 and E355 are critical for pH gating and cation selectivity (Schaub *et al*, 2020), and that the C-terminal leucine zipper may act in APOL1 dimerization (Schaub *et al*, 2021). Thus, the rod-like structure may contribute not only to oligomerization but also to enabling ion flux due to free access to the pore. However, this structure is observed only for G2, and not for APOL1 G1 or the wild type. It therefore remains unclear to what extent G1 and G2 allow ion flux through the pore via comparable mechanisms.

It remains to be shown to what extent APOL1’s ion conductance is enabled by missing intramolecular interactions, oligomerization of the RRVs, or interactions with other proteins. Future studies are also necessary to uncover how inter- or intramolecular interactions of APOL1 across the PM can be linked to the different clinical manifestations of AMKDs and used to develop therapeutic strategies.

## METHODS

### Constructs and cloning

For expression of APOL1 M1-G2 proteins, doxycycline (Dox) inducible plasmids in the pInducer21-puro expression system(Schulze *et al*, 2014) were generated using the Gateway™ cloning system. Base exchange of c.792C>A resulting in the amino acid variation p.N264K was achieved via site-directed mutagenesis using specific primers (fwd: gagaacatatccaaatttctttccttagc, rev: gctaaggaaagaaatttggatatgttctc) on existing pENTR-APOL1-G2 vectors. Mutated constructs were shuttled into pInducer21-puro expression vector using LR clonase™ enzyme mix II (11791100; Thermo Fisher). Amplification of constructs was performed in chemically competent *E. coli* and purification of plasmids was done using the Monarch® Plasmid miniprep Kit (T1010L; New England Biolabs) and the PureLinkTM HiPure Plasmid Maxiprep Kit (K210007; ThermoFisher Scientific). The untagged, C-terminal GFP and Glyc (Kaup *et al*, 2011) tagged plasmids of APOL1-G2 were already available and described elsewhere (Kaup *et al*, 2011; Granado *et al*, 2017; Müller *et al*, 2021). All used constructs possess the African haplotype “EIK”.

### Cell lines and cell culture

Cell lines were cultivated in Dulbecco’s modified eagle’s medium (DMEM, high glucose; D6429-500ML; Sigma Aldrich) with 10% FCS (FBS.S 0615; BIO & SELL) and 1% Penicillin-Streptomycin (P0781-100ML, Sigma Aldrich) at 37 °C and 5% CO_2_ atmosphere. Induction of overexpression in cells was achieved by adding 125 ng/ml doxycycline (Dox) to cultivation medium(Schulze *et al*, 2014; Granado *et al*, 2017; Müller *et al*, 2021).

HEK293T cells were stably modified with untagged and GFP tagged expression plasmids via lentiviral transduction. Virus generation was performed in HEK293T cells transfected via jetOPTIMUS® reagent (101000006, Satorius) with plasmid of choice and viral packaging plasmids before incubated for three days. Virus-containing supernatant was filtered (0.45 μm sterile filter), target HEK293T cells at a confluence of 60-70% were infected twice with a 1:1 ratio of virus suspension plus fresh medium containing polybrene (8 μg/ml). Infection cycles of 24 h were followed by regeneration phases of 24 h without virus. Afterwards, puromycin (4 μg/ml) was used for selection of cells. Degradation assay: To inhibit proteasomal activity, cells were treated with 5 µM MG-132 (M7449-1ML, Sigma-Aldrich) dissolved in DMSO for different time periods and analyzed by flow cytometry as described in detail earlier(Höffken *et al*, 2025a).

### Cell viability assays

Cell viability was detected in real time using the RealTime-Glo™ Cell Viability Assay (G9712, Promega) following the manufacturer’s instructions and as described before(Granado *et al*, 2017). Briefly, 10 000 HEK293T cells per well were seeded onto 96-well white bottom plates (Nunclon delta; 136101, ThermoFisher Scientific) in 50 µl cultivation medium. After 16-18 h adherence time, 2x RealTime-Glo™ enzyme-substrate mix (1:1) were diluted in 50 µl medium with or without 125 ng/ml Dox(Granado *et al*, 2017) and 50 ml mixture were added to each well. Luminescence was measured using a microplate reader (TECAN) at indicated time points. At least three independent experiments (N = 3) were measured for each cell line including six technical replicates (n = 6).

### Transient transfections

For transient expression of APOL1-GFP constructs, HEK293T cells at a confluency of 70-80% were transfected with jetOPTIMUS® (PolyPlus, 101000006) or Lipofectamine™2000 (Invitrogen, 11668500) reagents following the manufacturer’s instruction. For induction of overexpression 125 ng/ml doxycycline were simultaneously added to the medium. In brief, for jetOPTIMUS transfections 100 µl of buffer were mixed with 1 µg of DNA and 1 µl reagent for 1 ml medium volume per well, incubated for 10 min at RT and added to the cells. For Lipofectamine™2000 transfections 10 µl reagent and 4 µg DNA were each diluted in 250 µl OptiMEM (Gibco, 31985062) separately, incubated for 5 min, mixed with each other, incubated for 30 min at RT and added to the cells. After 6 h of transfection and induction, transfection medium was changed to cultivation medium containing Dox. After transfection and induction for 24 h, cells were washed with 1x phosphate-buffered saline (PBS) and processed for analysis.

### Preparation of cell lysates and PNGase F treatment

Cell lysate preparation and Western blotting analysis were performed as described earlier(Müller *et al*, 2021). Briefly, for preparation of lysates cells were grown on 12-well cell culture plates and washed with cold 1x PBS. For cell lysis, cells were transferred on ice and lysed in Lämmli buffer (20% (v/v) Glycerin, 125 mM Tris-HCl (pH 6.8), 10% (w/v) SDS, 0.2% (w/v) Brom phenole blue and 5% β-Mercaptoethanol in H_2_O) or for PNGase F treatment in RIPA buffer (50 mM Tris-HCl (pH 7.4), 150 mM NaCl, 1% NP-40, 0.5% Na-deoxycholate, 0.1% SDS) containing complete protease inhibitor (04693116001; Roche), phosphatase and protease inhibitor cocktails (P5726-5ML, P0044-5ML, P8340-1ML; Sigma Aldrich). To homogenize samples, lysates were passed ten times through a blunt 20-gauge needle (0.9 mm diameter), RIPA lysates were additionally incubated & vortexed on ice for 30 min before sonicated for 10 min. Lämmli lysates were denatured by heating for 10 min at 95 °C, while RIPA lysates were centrifuged (20-30 min, 4 °C, 14 000 x g) to remove cell debris. RIPA supernatants and Lämmli lysates were stored at −20 °C for further use. Glycosylation status was investigated via digestion of N-linked oligosaccharides in RIPA lysates with PNGase F (P0704L, New England Biolabs) according to the manufacturer’s instructions. Digested samples were analyzed by Western blot.

### Western blot analyses

Wetsern blot were perforemeed as described earlier (Schulze *et al*, 2014; Granado *et al*, 2017; Müller *et al*, 2021). In brief, sample separation was achieved by 10% SDS PAGEs (200 V, 45 min). RIPA lysates were mixed with 2× Lämmli buffer and proteins were denatured (95 °C, 5 min) before loaded to the SDS PAGE. Proteins were semi-dry blotted onto methanol activated PVDF membranes (0.45 μm pore size; IPVH00010; Merck Millipore, Burlington, MA, USA) for 90 min at 1 mA/cm^3^. After transfer, membranes were blocked with 5% milk blocking (T145.2, Roth) in TBST for 30 min and incubated with rabbit anti-APOL1 antibody (1:1000; HPA0018885, Sigma) or rabbit anti-α-Actinin4 antibody (1:1000; S15145S, Cell Signaling) diluted in 5% bovine serum albumin (BSA; 8076.3, Roth) in TBST overnight at 4 °C. After washing membranes three times in TBST, secondary horseradish peroxidase-coupled goat anti-rabbit IgG antibody (1:2000; 115-035-144; Jackson ImmunoResearch) was diluted in blocking solution and added to membranes for 1 h at RT. Unbound antibodies were removed by washing with TBST three times and proteins were detected with Clarity™ Western ECL substrate (1705061; BioRad) on an Azure c600 Compare cSeries imaging system (Azure Biosystems Inc., Dublin, CA, USA).

### Immunofluorescence (IF) microscopy

Staining of APOL1 at the plasma membrane was performed similar as described elsewhere(Gupta *et al*, 2020). In brief, after 24 h of induction cells on coverslips were fixed for 15 min in 4% paraformaldehyde (PFA; P087.1, Roth). After carefully washing twice with 1x PBS for 10 min, unpermeabilized cells were incubated for 1 h on ice with primary rabbit anti-APOL1 antibody (11486-2-AP, ProteinTech) diluted 1:200 in PBS. After washing twice in PBS, cells were incubated with secondary fluorophore-coupled goat anti-rabbit-AlexaFluor®647 antibody (A-21244, Invitrogen) diluted 1:1500 in PBS for 1 h on ice. Counterstaining of DNA was performed with DAPI (1:5000) for 5 min before cells were rinsed in tap water and mounted with Mowiol. HEK293T wt cells incubated with the APOL1-specific antibody as well as APOL1-G2-GFP induced cells lacking incubation with the primary antibody served as controls (Suppl. Fig. SF1). Samples were analyzed using a fluorescence microscope (Axio Observer Z1, HXP120, Axiocam MRm; Carl Zeiss) with EC Plan-Neofluar 63x objective using immersion oil (Zeiss) and corresponding Zen Blue software (v2.3).

### Flow cytometry (FC)

For FC analysis, cells were grown and treated on 12-well cultivation plates according to protocols descried before(Gupta *et al*, 2024; Höffken *et al*, 2025a). In brief, cells were washed twice with 1x PBS, resolved in 500 µl PBS and transferred into a 1.5 ml microreaction tube. For fixation, 500 µl 4% PFA were added, and cells were fixed in this 2% PFA/PBS mixture for 15 min at RT. After centrifugation for 5 min at 1000 x g and 4 °C supernatant was discarded, cells were resuspended in PBS and stored dark at 4 °C until further processing or measurement. Staining of the APOL1 surface level was performed similarly as for microscopy and described before(Gupta *et al*, 2020). After washing fixed cells twice with 2 ml cold PBS and centrifugation, cells were resuspended and incubated for 1 h on ice in 100 µl primary rabbit anti-APOL1 antibody (11486-2-AP, ProteinTech) diluted 1:200 in PBS. After a double washing and centrifugation step, cells were incubated in 100 µl secondary fluorophore-coupled goat anti-rabbit-AlexaFluor®647 antibody (AF647, A-21244, Invitrogen) diluted 1:1500 in PBS for 1 h on ice. Finally, cells were washed with PBS, resuspended in 400 µl PBS and stored at 4 °C in the dark until measurement.

FC cytometry analysis was performed using a FACSCalibur™ Flow cytometer (BD Biosciences) with the corresponding CellQuest Pro software using a blue argon laser (488 nm) and a red diode laser (635 nm). Cell size and internal complexity (e.g. granularity) were detected using forward scatter (FSC) and sideward scatter (SSC) detectors, respectively. Fluorescence signals were detected according to GFP signal (515-545 nm) or AF647 signal (653-669 nm). For measurement of GFP and AF647 intensities of cells with stained surface APOL1, ≥ 50 000 viable cells were measured.

For FC data analysis FlowJo (v10.10.0) was used. Cells were gated for viable cell population with a FSC height >150. As especially the APOL1-G2 cell lines contained a subpopulation which seemed not inducible anymore, those were gated using the GFP signal of HEK293T wt cells and excluded as wt-like signals if necessary. Remaining populations were defined as APOL1-positive inducible subpopulations. Subsequently, the GFP negative signal for each stable cell line was defined using the 95^th^ percentile of GFP intensity of uninduced viable cells (-Dox). Unspecific signals of surface APOL1 staining were gated by the 95^th^ percentile of AF647 intensity of viable HEK293T wildtype control. For surface stained APOL1, specific detection of secondary antibody was tested using omission control, lacking incubation with the primary antibody. For transient transfections, GFP and AF647 intensities were gated using untransfected HEK293T cells as ctr. Combined gates for GFP^+^ and AF647^+^ signals account for positively transfected cells which exhibit PM APOL1 signal (Q2).

### *In silico* APOL1 pore modeling

The part of the APOL1 amino acid sequence that included the previously predicted transmembrane (TM) region (aa 171-360) was modelled using *AlphaFold 3*(Abramson *et al*, 2024). The simulation was performed by adding myristic acid and sodium ions; all other parameters were set to default values. The threshold for the predicted template modeling (pTM) score was set at 0.5. The C-terminal part (aa 171-396) of the APOL1-G2 was modelled with AlphaFold3 following the same protocol as described above.

The alternative tool *RosettaFold*(Baek *et al*, 2021), a deep learning-based method integrated into the protein structure prediction server *Robetta*, was used to verify prediction and lead to similar results. As *RosettaFold* did not include fatty acids as membranous surrounding of the pore, AlphaFold3 was chosen for further modeling (Suppl. Fig. SF2).

The predicted *AlphaFold3* model, including the four TM regions, was prepared at pH 7.5 ± 0.5 to assign appropriate protonation states(Johnston *et al*, 2023). Then, it was inserted into a Desmond default POPC membrane bilayer, and the system was solvated. 0.15 M NaCl, KCl, or CaCl_2_ was added to the system. Schrödinger Desmond(Bowers *et al*, 2006) was used to perform Molecular Dynamics (MD) simulations. The simulation time was set to 200 ns, and the NPAT ensemble class was used, with a temperature of 300 K and a pressure of 1.013 bar. After MD simulations, the trajectories were analyzed, and the data were used to plot different graphs. The SSE plot shows the evolution of protein secondary structure elements throughout the simulation. It reports the SSE distribution by residue index throughout the protein structure, with residue 1 corresponding to the p.171 of APOL1. The RMSD graph displays the evolution of the protein throughout the simulation. All frames were aligned on the reference frame backbone, and then the RMSD was calculated (Suppl. Fig. SF2).

To analyze the M1 (N264K) variant, TM-based structure of APOL1 was subjected to *in silico* mutagenesis to generate the p.N264K variant. After protein preparation, the MD simulation was carried out as described above, using NaCl as a salt to neutralize the system. The model of C-terminal part of the G2 variant of APOL1 was used to predict multimeric assembly with the ClusPro server for protein-protein docking(Jones *et al*, 2022).

### Statistical analyses and figure preparation

Statistical analysis of viability data was performed using 2-way ANOVA with Šídák’s multiple comparisons test and FACS data were analyzed using RM 1-way ANOVA with Dunnett’s test for multiple comparison or paired *t*-test for transient transfections. Significance was calculated between uninduced condition (-Dox) and treated samples or between ctr and induced cells for transient transfection data. Levels of significance of an adjusted p value are indicated by asterisks: ^*^ p ≤ 0.05, ^**^ p ≤ 0.01, ^***^ p ≤ 0.001 and ^****^ p ≤ 0.0001 and ns for non significant. Data are presented as means with SEM. Graphs and statistics were generated using GraphPad Prism (v10.6.1). Images were further processed using Zen Blue (v2.3) and ImageJ/FIJI. Figures were prepared using CorelDraw2024. Schematic illustrations were generated using BioRender (https://BioRender.com).

## Supporting information

uncropped_western_blots

## ACKNOWLEDGMENTS

We thank the DFG for funding, all members of the laboratory for their valuable discussions, and Dr. Giuliano Ciarimboli for kindly bringing Italian and German researchers together and for critical reading the manuscript.

## AUTHOR CONTRIBUTIONS

The conceptualization of the project was carried out by VH, HP, CI, and TW. Cell biological experiments were performed by VH and NN. The 3D analyses were conducted by LC and CI. Supervision was provided by VH, IC, HP, and TW. Writing, review and editing were done by VH, LC, and TW. Funding acquisition was undertaken by TW.

## THE PAPER EXPLAINED

PROBLEM: Certain genetic variants of the APOL1 gene greatly increase the risk of kidney disease, especially among people of African ancestry. Our study investigates a newly discovered variant, called p.N264K (M1), which appears to protect against this risk in individuals carrying the APOL1-G2 variant.

RESULTS: The protective p.N264K variant does not change APOL1’s expression, localization, or turnover in cells. Instead, our structural and computational analyses show that the lysine at position 264 acts like a plug inside the protein’s transmembrane pore, impairing monovalent cation binding and perhaps ion flux. In addition, *in silico* modeling suggests that the G2 form of APOL1 can assemble into multimeric structures that may enable pore activity.

IMPACT: These findings reveal a clear structural mechanism behind the protective effect of M1 and open new possibilities for developing drugs that mimic the pore-blocking action to prevent or treat APOL1-related kidney disease.

## DATA AVAILABILITY STATEMENT

The data that support the findings of this study are available from the corresponding author upon reasonable request. Supplementary Material is available online.

## FIGURES and FIGURE LEGENDS

**Expanded View Figure EV1:**
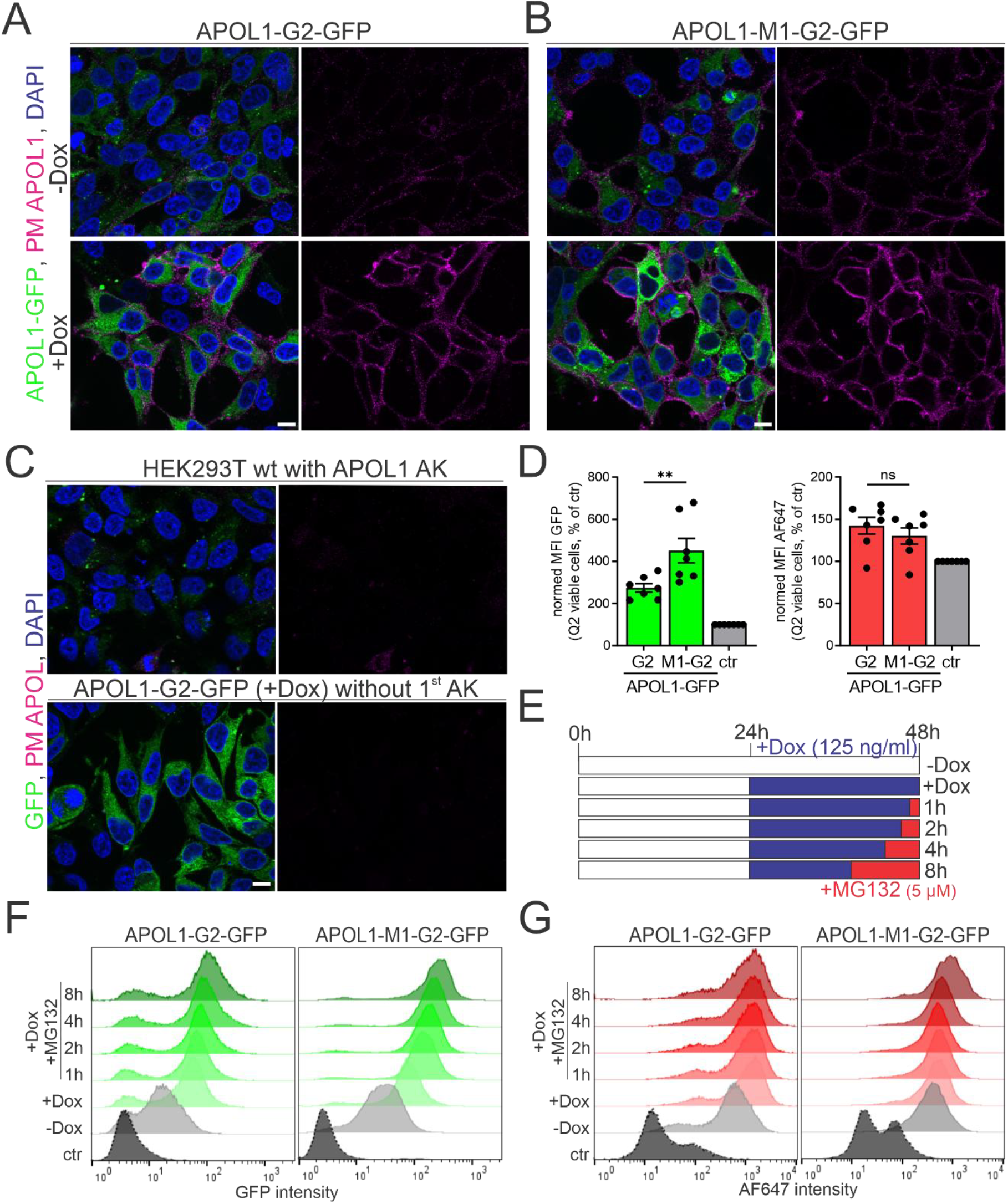
Surface-localized APOL1 pools were detected in both the M1 variant and APOL1-G2 indicating efficient transport to the PM. **(A, B)** Immunofluorescence (IF) staining of unpermeabilized HEK293T cells expressing APOL1-G2-GFP or with additional M1 variant in uninduced cells (-Dox) and after 24 h of Dox induction (+Dox, 125 ng/ml). GFP signals display total APOL1 protein within the cells, specific antibody staining against APOL1 (magenta) stained surface-localized APOL1 proteins and nuclei were counterstained with DAPI. Scalebars = 10 µm. **(C)** HEK293T wt cells incubated with the specific APOL1 antibody and Dox-induced APOL1-G2-GFP cells lacking incubation with primary antibody were used as controls for specificity IF staining **(D)** For a direct comparison of APOL1-G2 and APOL1-M1-G2, HEK293T cells were transiently transfected with equal amounts of the same expression plasmids (only the cDNA differs) and induced with Dox for 24 h (125 ng/ml). Normalized mean fluorescence intensities (MFI) of GFP (left) and AF647 (right) in these cells showed positive expression of APOL1-GFP for both constructs by flow cytometry, but higher in M1-G2, as well as a similar presence of surface-localized pools for both the G2 and M1-G2 variants. (N = 7) ^**^/ns = 0.001/not significant. Values are presented as means ± SEM. (E) Experimental scheme of degradation assay: HEK293T cells expressing C-terminally GFP-tagged APOL1 forms G2 and M1-G2 were induced for 24 h (+Dox, 125 ng/ml) with or without MG132 treatment (5 µM) for indicated timepoints. **(F, G)** Exemplary histograms of fluorescence intensities of GFP **(E)** representing total APOL1 and AF647 (F) displaying PM-APOL1 show proportional increase of GFP signals with inhibition of degradation for both variants while PM APOL1 signals remained more stable.

**Expanded View Figure EV2:**
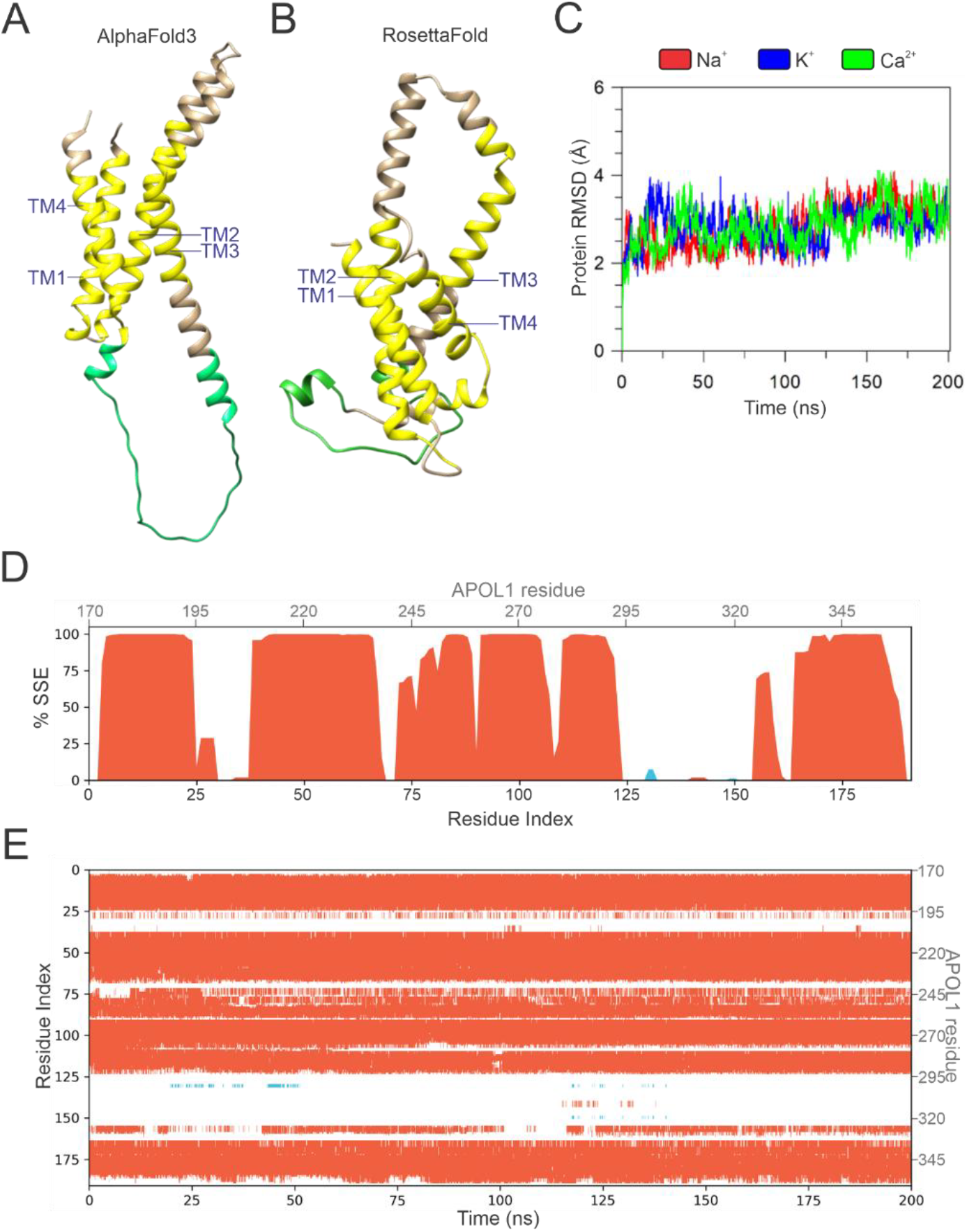
In silico prediction and validation of the APOL1 TM regions (aa 171-360). **(A, B)** Structures of the transmembrane segments (aa 171-360) of APOL1 predicted by *AlphaFold3* (A) and *RobettaFold* (B) verified presence of four helices as putataive transmembrane regions (yellow). Predictions are similar but show different orientations of TM3 and the following loop (green). **(C)** The plots illustrate the RMSD evolution of the modeled TM structure during the MD simulation of APOL1-G0 in the presence of sodium (red), potassium (blue), or calcium (green). **(D, E)** Protein secondary structure elements (SSE), such as α-helices, were monitored and persist throughout the simulation. Shown is the distribution of SSEs by residue index across the used sequence, with the first residue corresponding to p.171 of the APOL1 sequence over the simulation time.

## REFERENCES

Abramson J, Adler J, Dunger J, Evans R, Green T, Pritzel A, Ronneberger O, Willmore L, Ballard AJ, Bambrick J, et al (2024) Accurate structure prediction of biomolecular interactions with AlphaFold 3. Nature 630: 493– 500

Baek M, DiMaio F, Anishchenko I, Dauparas J, Ovchinnikov S, Lee GR, Wang J, Cong Q, Kinch LN, Dustin Schaeffer R, et al (2021) Accurate prediction of protein structures and interactions using a three-track neural network. Science (80-) 373

Bowers KJ, Chow E, Xu H, Dror RO, Eastwood MP, Gregersen BA, Klepeis JL, Kolossvary I, Moraes MA, Sacerdoti FD, et al (2006) Scalable algorithms for molecular dynamics simulations on commodity clusters. In Proceedings of the 2006 ACM/IEEE Conference on Supercomputing, SC’06

Bruno J, Pozzi N, Oliva J & Edwards JC (2017) Apolipoprotein L1 confers pH-switchable ion permeability to phospholipid vesicles. J Biol Chem 292: 18344–18353

Cuypers B, Lecordier L, Meehan CJ, Van den Broeck F, Imamura H, Büscher P, Dujardin J-C, Laukens K, Schnaufer A, Dewar C, et al (2016) Apolipoprotein L1 Variant Associated with Increased Susceptibility to Trypanosome Infection. MBio 7

Daneshpajouhnejad P, Kopp JB, Winkler CA & Rosenberg AZ (2022) The evolving story of apolipoprotein L1 nephropathy: the end of the beginning. Nat Rev Nephrol 18 doi:10.1038/s41581-022-00538-3 [PREPRINT]

Datta S, Antonio BM, Zahler NH, Theile JW, Krafte D, Zhang H, Rosenberg PB, Chaves AB, Muoio DM, Zhang G, et al (2024) APOL1-mediated monovalent cation transport contributes to APOL1-mediated podocytopathy in kidney disease. J Clin Invest 134

Friedman DJ & Pollak MR (2021) Apol1 nephropathy: From genetics to clinical applications. Clin J Am Soc Nephrol 16

Genovese G, Friedman DJ, Ross MD, Lecordier L, Uzureau P, Freedman BI, Bowden DW, Langefeld CD, Oleksyk TK, Uscinski Knob AL, et al (2010) Association of Trypanolytic ApoL1 Variants with Kidney Disease in African Americans. Science (80-) 329: 841–845

Giovinazzo JA, Thomson RP, Khalizova N, Zager P, Malani N, Rodriguez-Boulan E, Raper J & Schreiner R (2020) Apolipoprotein L-1 renal risk variants form active channels at the plasma membrane driving cytotoxicity. Elife

Granado D, Müller D, Krausel V, Kruzel-Davila E, Schuberth C, Eschborn M, Wedlich-Söldner R, Skorecki K, Pavenstädt H, Michgehl U, et al (2017) Intracellular APOL1 Risk Variants Cause Cytotoxicity Accompanied by Energy Depletion. J Am Soc Nephrol

Gupta N, Waas B, Austin D, De Mazière AM, Kujala P, Stockwell AD, Li T, Yaspan BL, Klumperman J & Scales SJ (2024) Apolipoprotein L1 (APOL1) renal risk variant-mediated podocyte cytotoxicity depends on African haplotype and surface expression. Sci Rep 14

Gupta N, Wang X, Wen X, Moran P, Paluch M, Hass PE, Heidersbach A, Haley B, Kirchhofer D, Brezski RJ, et al (2020) Domain-Specific Antibodies Reveal Differences in the Membrane Topologies of Apolipoprotein L1 in Serum and Podocytes. J Am Soc Nephrol

Gupta Y, Friedman DJ, McNulty MT, Khan A, Lane B, Wang C, Ke J, Jin G, Wooden B, Knob AL, et al (2023) Strong protective effect of the APOL1 p.N264K variant against G2-associated focal segmental glomerulosclerosis and kidney disease. Nat Commun 14

Höffken V, Alvermann L, Niggemeier D, Beul K, Nedvetsky P, Ellinger B, Assenmacher D, Granado D, Pavenstädt H & Weide T (2025a) Renal disease factor APOL1: plasma membrane pools resist rapid turnover.

Höffken V, Braun DA, Pavenstädt H & Weide T (2025b) A Cell Biologist’s View on APOL1: What We Know and What We Still Need to Address.

Hung AM, Assimon VA, Chen HC, Yu Z, Vlasschaert C, Triozzi JL, Chan H, Wheless L, Wilson O, Shah SC, et al (2023) Genetic Inhibition of APOL1 Pore-Forming Function Prevents APOL1-Mediated Kidney Disease. J Am Soc Nephrol 34

Johnston RC, Yao K, Kaplan Z, Chelliah M, Leswing K, Seekins S, Watts S, Calkins D, Chief Elk J, Jerome S V., et al (2023) Epik: pKa and Protonation State Prediction through Machine Learning. J Chem Theory Comput 19

Jones G, Jindal A, Ghani U, Kotelnikov S, Egbert M, Hashemi N, Vajda S, Padhorny D & Kozakov D (2022) Elucidation of protein function using computational docking and hotspot analysis by ClusPro and FTMap. Acta Crystallogr Sect D Struct Biol 78

Kaup M, Saul VV, Lusch A, Dörsing J, Blanchard V, Tauber R & Berger M (2011) Construction and analysis of a novel peptide tag containing an unnatural N-glycosylation site. FEBS Lett

Lecordier L, Heo P, Graversen JH, Hennig D, Skytthe MK, Cornet d’Elzius A, Pincet F, Pérez-Morga D & Pays E (2023) Apolipoproteins L1 and L3 control mitochondrial membrane dynamics. Cell Rep 42

Müller D, Schmitz J, Fischer K, Granado D, Groh A-C, Krausel V, Lüttgenau SM, Amelung TM, Pavenstädt H & Weide T (2021) Evolution of Renal-Disease Factor APOL1 Results in Cis and Trans Orientations at the Endoplasmic Reticulum That Both Show Cytotoxic Effects. Mol Biol Evol

Olabisi OA, Zhang J-Y, VerPlank L, Zahler N, DiBartolo S, Heneghan JF, Schlöndorff JS, Suh JH, Yan P, Alper SL, et al (2016) APOL1 kidney disease risk variants cause cytotoxicity by depleting cellular potassium and inducing stress-activated protein kinases. Proc Natl Acad Sci U S A 113: 830–7

Pays E, Radwanska M & Magez S (2023) The Pathogenesis of African Trypanosomiasis. Annu Rev Pathol Mech Dis 18 doi:10.1146/annurev-pathmechdis-031621-025153 [PREPRINT]

Pays E, Vanhollebeke B, Vanhamme L, Paturiaux-Hanocq F, Nolan DP & Perez-Morga D (2006) The trypanolytic factor of human serum. NatRevMicrobiol 4: 477–486

Perez-Morga D, Vanhollebeke B, Paturiaux-Hanocq F, Nolan DP, Lins L, Homblé F, Vanhamme L, Tebabi P, Pays A, Poelvoorde P, et al (2005) Microbiology: Apolipoprotein L-I promotes trypanosome lysis by forming pores in lysosomal membranes. Science (80-) 309: 469–472

Schaub C, Lee P, Racho-Jansen A, Giovinazzo J, Terra N, Raper J & Thomson R (2021) Coiled-coil binding of the leucine zipper domains of APOL1 is necessary for the open cation channel conformation. J Biol Chem 297

Schaub C, Verdi J, Lee P, Terra N, Limon G, Raper J & Thomson R (2020) Cation channel conductance and pH gating of the innate immunity factor APOL1 are governed by pore-lining residues within the C-terminal domain. J Biol Chem doi:10.1074/jbc.RA120.014201 [PREPRINT]

Schulze U, Vollenbröker B, Braun DADADA, Van Le T, Granado D, Kremerskothen J, Fränzel B, Klosowski R, Barth J, Fufezan C, et al (2014) The Vac14-interaction Network Is Linked to Regulators of the Endolysosomal and Autophagic Pathway. Mol Cell Proteomics 13

Simeone CA, McNulty MT, Gupta Y, Genovese G, Sampson MG, Sanna-Cherchi S, Friedman DJ & Pollak MR (2025) The APOL1 p.N264K variant is co-inherited with the G2 kidney disease risk variant through a proximity recombination event. G3 Genes, Genomes, Genet 15

Tzur S, Rosset S, Shemer R, Yudkovsky G, Selig S, Tarekegn A, Bekele E, Bradman N, Wasser WG, Behar DM, et al (2010) Missense mutations in the APOL1 gene are highly associated with end stage kidney disease risk previously attributed to the MYH9 gene. Hum Genet 128: 345–350

Vanhamme L, Paturiaux-Hanocq F, Poelvoorde P, Nolan DP, Lins L, Van Den Abbeele J, Pays A, Tebabi P, Van Xong H, Jacquet A, et al (2003) Apolipoprotein L-I is the trypanosome lytic factor of human serum. Nature 422: 83–7

