## Supplementary material for "The APOL1 variant p.N264K blocks ion flow by occluding a pore at the cell surface": uncropped_western_blots

*by*

Verena Hoeffken[1*], Lara Console[2*], Niklas Nelde[1], Hermann Pavenstädt[1], Cesare Indiveri[2§], and Thomas Weide[1§]

Uncropped Western blot images

Figure 1: Inducible expression of the M1 variant of APOL1-G2 showed no cytotoxicity while similar orientation within the ER membrane.

Fig. 1 B)

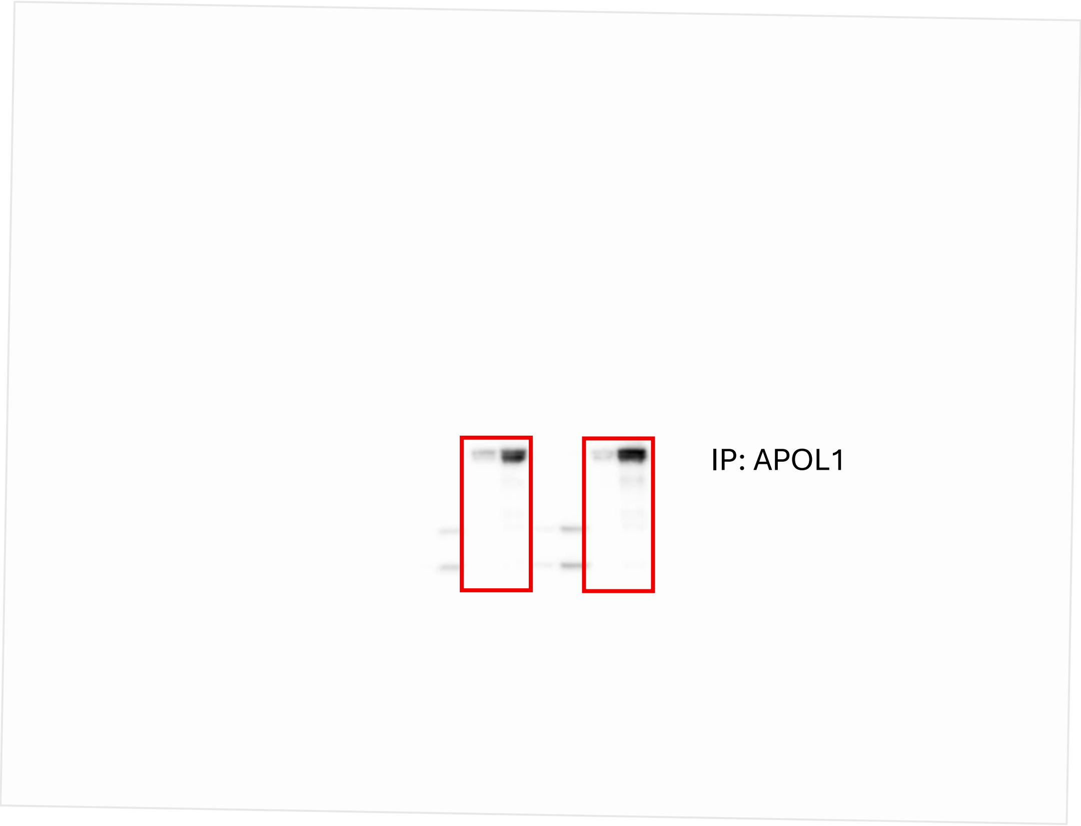

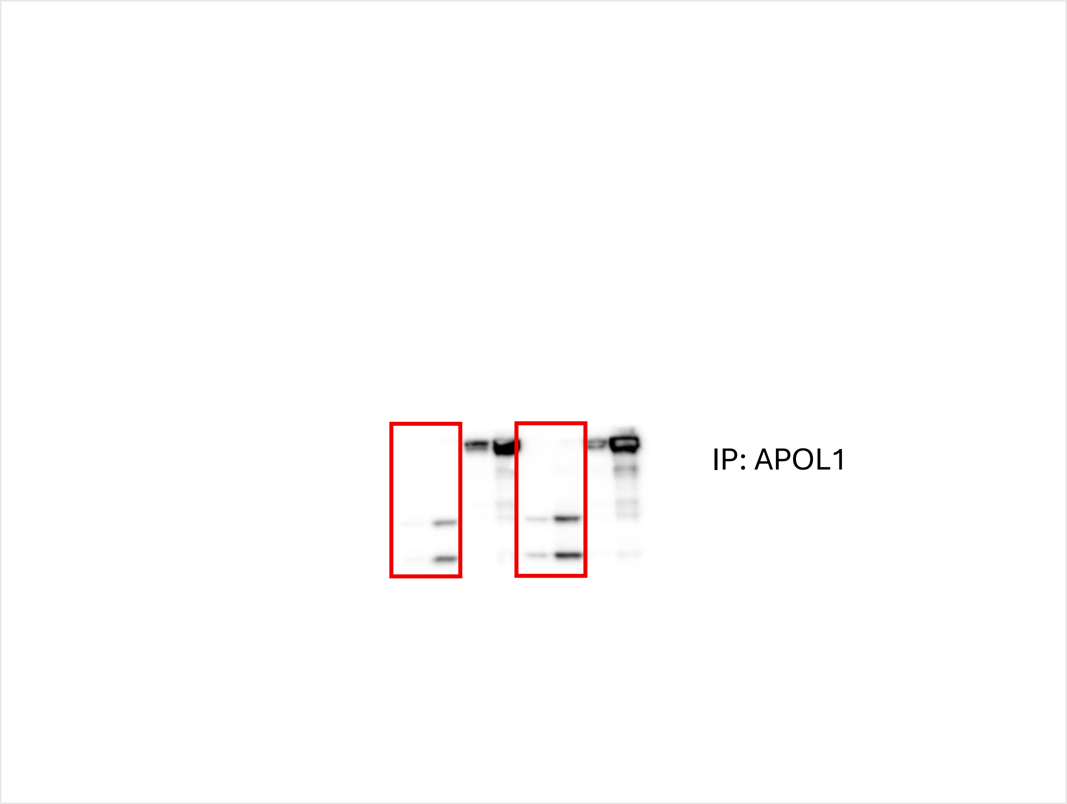


16

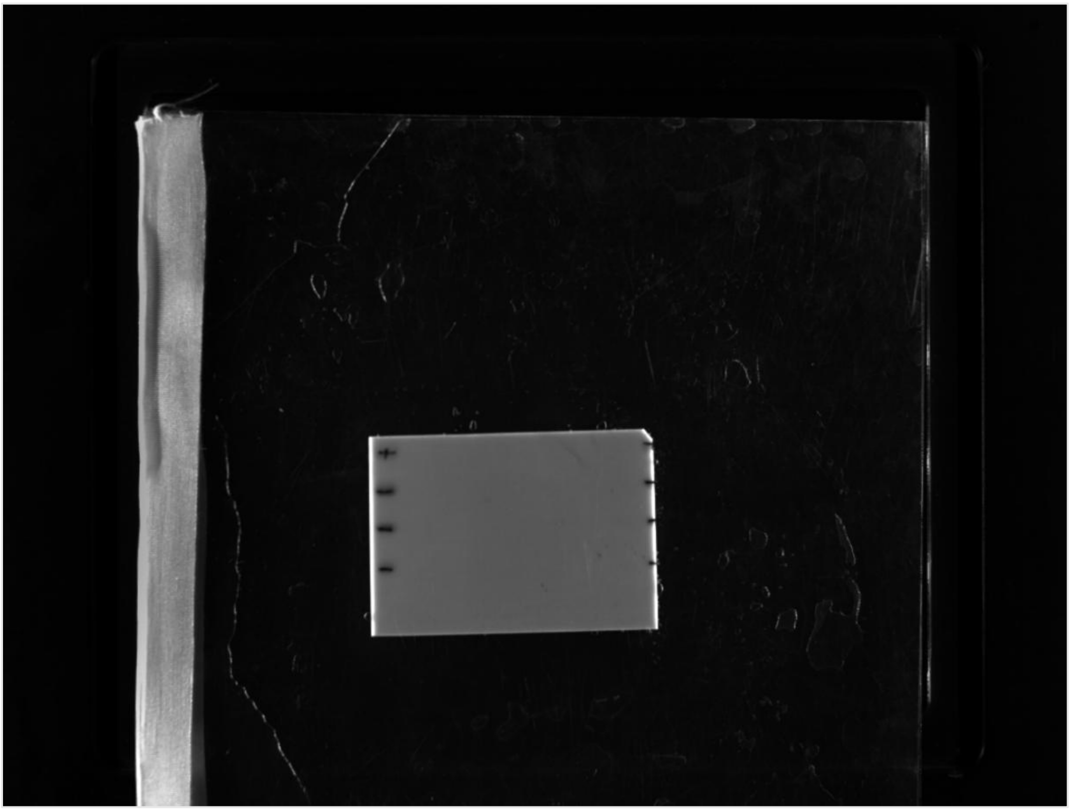

17

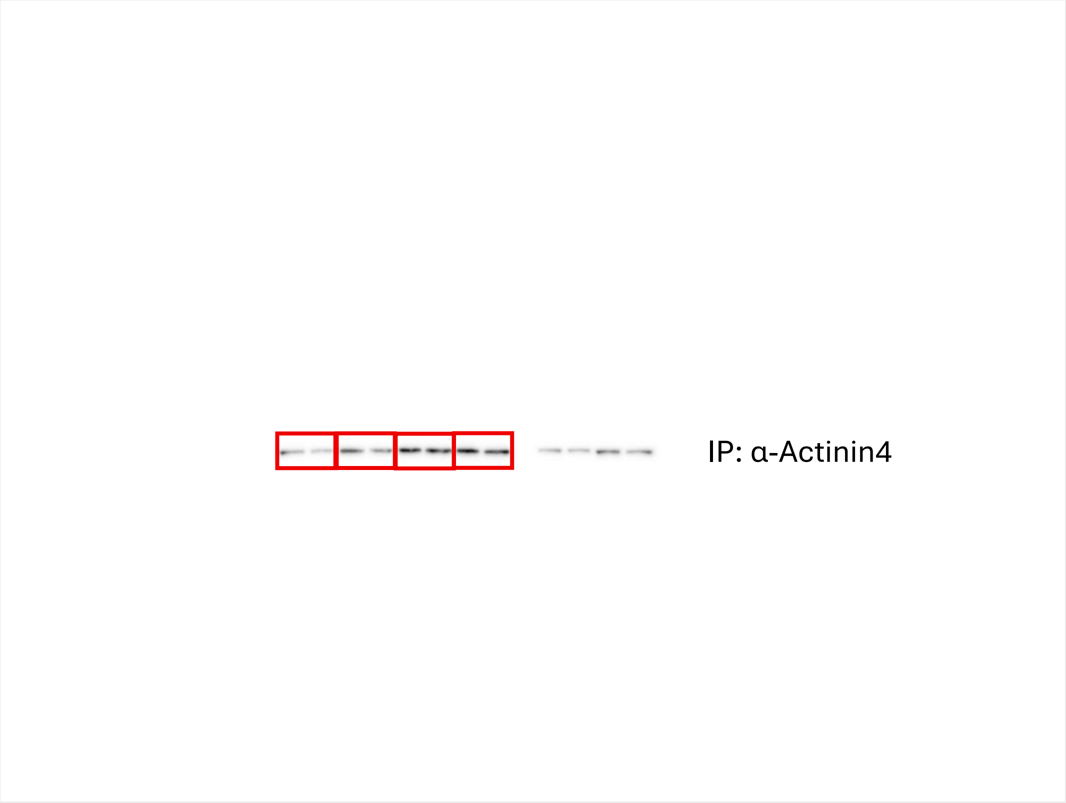


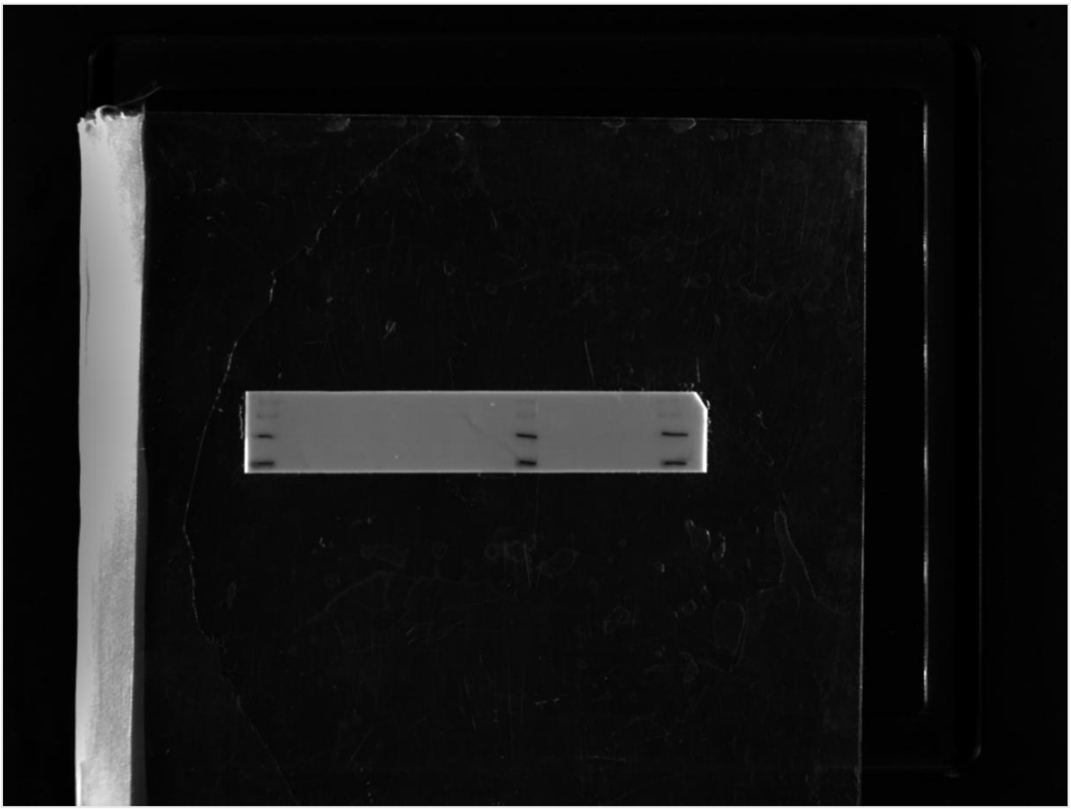

Fig. 1 E)

22

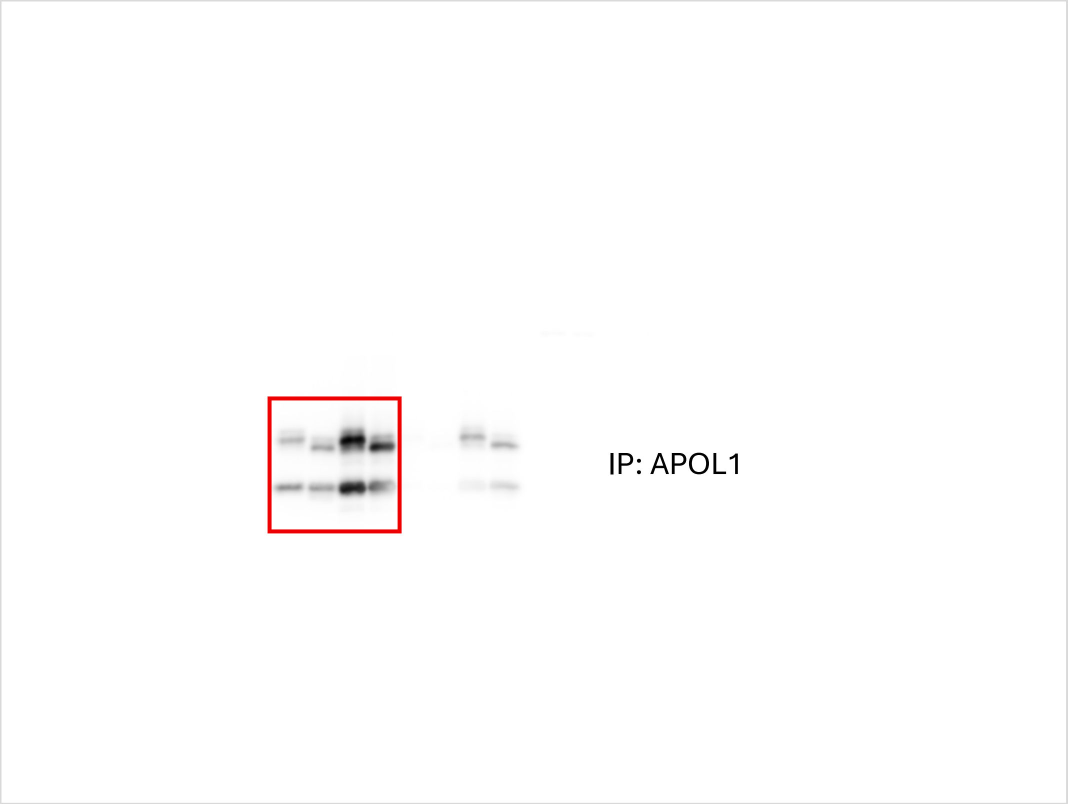


23

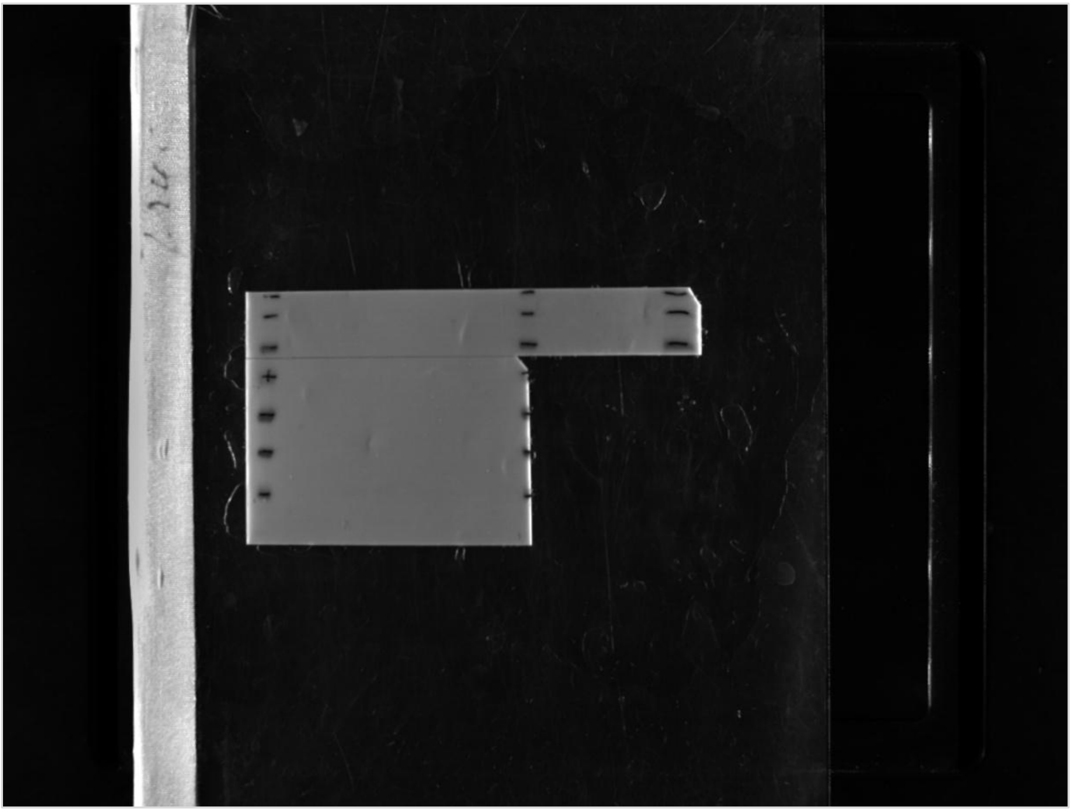

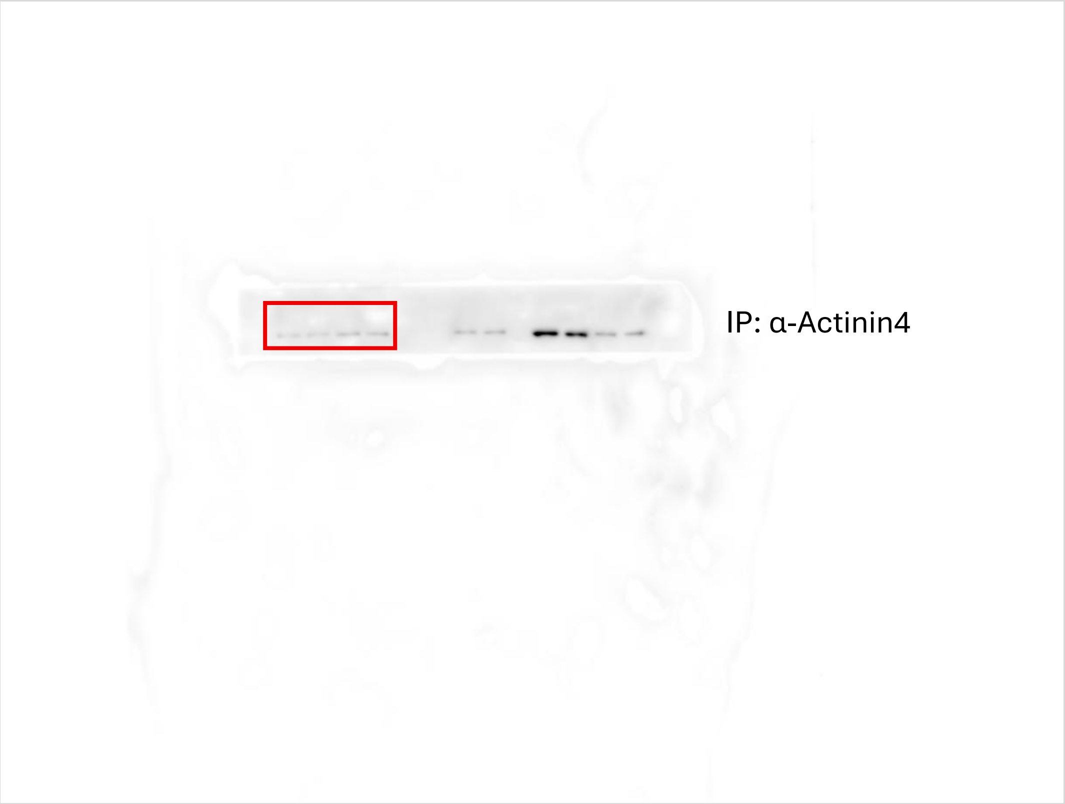
IP: α-Actinin4

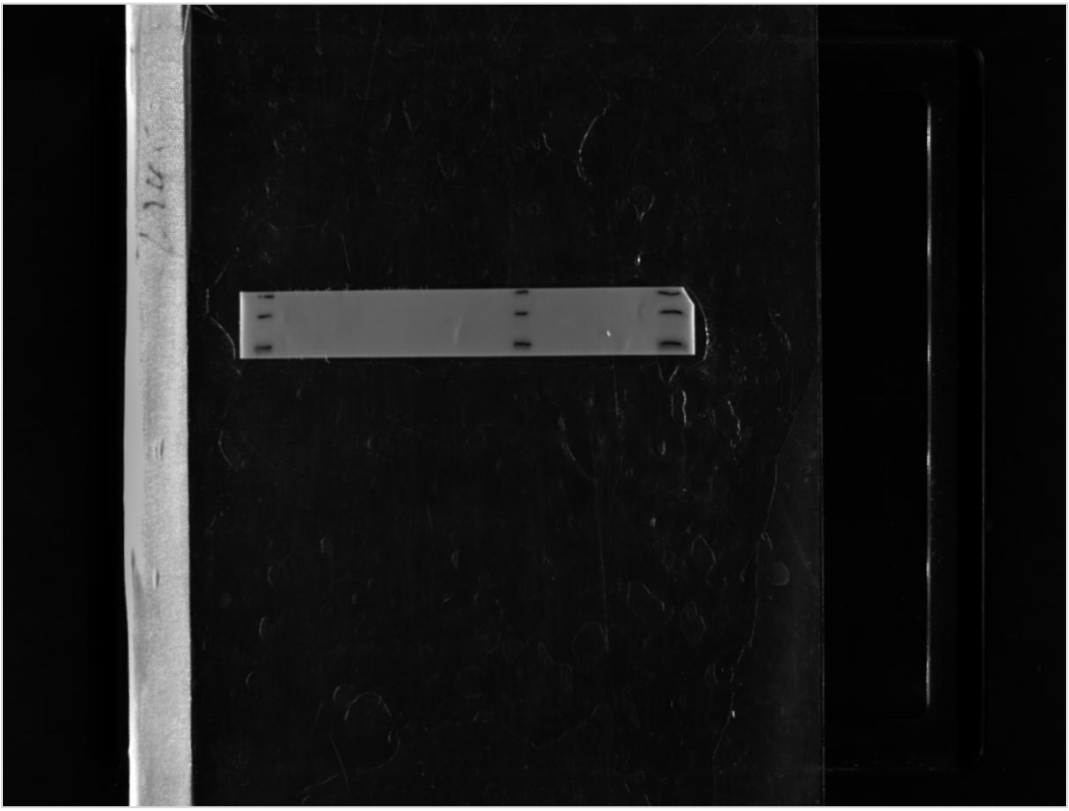
